# Combined genomic and molecular analysis defines prognostic markers of relapse in stage IA-IC1 clear cell ovarian carcinoma

**DOI:** 10.1101/2025.03.20.644313

**Authors:** Yasushi Iida, Michael Churchman, Robert L Hollis, Sarah Taylor, Clare Bartos, Ian Croy, William Gentleman, Tzyvia Rye, Rachel Nirsimloo, C. Simon Herrington, Charlie Gourley, Aikou Okamoto, John P Thomson

## Abstract

**Objective:** Clear cell ovarian carcinoma (CCOC) is generally associated with a favourable prognosis, but up to 30% of low-stage cases relapse within five years. The benefit of adjuvant chemotherapy for low-stage CCOC (FIGO stage IA-IC1) remains uncertain. This study aimed to identify molecular and immune markers associated with relapse in a well-characterized CCOC cohort.

**Methods:** Using the Edinburgh Ovarian Cancer Database, we identified 85 CCOC cases. Targeted DNA sequencing assessed genomic alterations and tumour immune infiltration was evaluated using CD3 and CD8 immunohistochemistry on tissue microarrays. Tumours were stratified by stage (IA-IC1, IC2-II, III-IV), and both univariate and multivariate analysis applied for progression-free survival (PFS) included stage, age, genomic features, and immune markers.

**Results:** Common genomic alterations included *ARID1A* (49%) and *PIK3CA* (42%) mutations, PIK3-AKT pathway perturbations (60%), and mismatch repair-related mutational signatures (25.9%). Genomic and molecular features were not significantly associated with tumour stage; however, low-stage tumours (IA-IC1) were enriched in CD3+ (40% cases) and CD8+ (29% cases) tumour-infiltrating lymphocytes (TILs) compared to higher-stage tumours (IC2-II: 13%/7%; III-IV: 9%/0% cases). In univariate analysis, low CD3+ TIL levels were significantly associated with reduced progression-free survival (HR = 4.4, P = 0.042, 95% CI 1.1-18). *ARID1A* wild-type status was linked to poorer PFS but only in low-stage tumours HR = 7.2, P = 0.088; 95% CI 0.74-72). Notably, the combination of *ARID1A* wild-type status and CD3+ depletion identified a high-risk subgroup among low-stage CCOC cases (HR = 11.7, P = 0.050; 95% CI 1-138), highlighting potential as a composite risk marker set.

**Conclusions:** Combined *ARID1A* mutation and low CD3+ve TIL levels are suggestive of higher recurrence risk in low-stage CCOC patients. These findings support further investigation of targeted therapies or immune checkpoint inhibitors in larger CCOC cohorts.

**Highlights:**

- We report a well-characterized CCOC cohort with rich clinical, genomic and molecular data.
- Common mutations included *ARID1A* (49%) and *PIK3CA* (42%), with MSI in 9%, showing no significant enrichment across tumour stages.
- Low-stage CCOC has higher levels of CD3+ & CD8+ tumour infiltrating leukocytes than in advanced-stage disease.
- ARID1A wild-type tumours with CD3+ TIL depletion represent a high-risk subgroup of low-stage CCOC.
- Findings support further study and highlight potential markers for recurrence risk.

## Introduction

Clear cell ovarian carcinoma (CCOC) is a rare but distinct subtype of epithelial ovarian cancer, accounting for approximately 5–10% of all cases [1]. It is characterized by unique clinical and molecular features, including a relatively high prevalence of low-stage diagnoses, particularly in International Federation of Obstetrics and Gynaecology FIGO stage IA-IC1 tumours. While low-stage CCOC generally carries a more favourable prognosis compared to advanced-stage disease, a significant proportion of patients relapse within five years of diagnosis, leading to poor survival outcomes [1]. These relapses highlight a critical gap in our ability to identify high-risk patients, particularly within low-stage cohorts, and underscore the need for robust prognostic biomarkers.

Previous studies have highlighted that somatic mutations in the SWI/SNF complex are common, particularly in *ARID1A* (49-67%); as well as those in the PI3K/AKT/mTOR pathway, including *PIK3CA* (42-54%), and the RTK/RAS pathway, including *KRAS* (13-15%) [2-8]. Despite their prevalence, the prognostic implications of these mutations, particularly in relation to relapse risk and survival, remain poorly understood.

Emerging evidence also points to the importance of the tumour immune microenvironment in determining patient outcomes. Immune cell infiltration, particularly by CD3+ T cells and CD8+ cytotoxic T cells, has been shown to influence survival across various cancers, including ovarian cancer [9-11]. Recent efforts to characterize the immune microenvironment in CCOC have found no correlation between overall survival and immune infiltration in treatment-naïve CCOC; however, CD8+ T cell expression levels were found to increase upon recurrence [12]. Beyond this, the role of immune infiltration in CCOC and how this is associated with specific genomic alterations, warrants investigation.

Currently, the management of low-stage CCOC, comprising FIGO stages IA-IC1, is challenging. Unlike high-grade serous ovarian carcinoma (HGSOC), where adjuvant chemotherapy is the standard of care, the benefit of chemotherapy in low-stage CCOC remains unclear [1, 13]. This uncertainty is compounded by the distinct biological characteristics of CCOC, which include limited responsiveness to platinum-based chemotherapy and unique genomic and immune profiles [1].

We therefore sought to identify novel biomarkers associated with relapse and survival in CCOC, particularly when stratified by disease stage. Through robust pathological review follow by targeted DNA sequencing and immune cell infiltration analysis, we set out to define molecular and immunological features that stratify patients based on their risk of recurrence. Specifically, our study aimed to identify prognostic markers that could guide clinical decision-making and improve treatment stratification. Additionally, we explore whether combining genomic features, such as *ARID1A* mutational status, with measures of immune infiltration could enhance prognostic precision in the low stage setting with a view towards identifying opportunities for personalized therapeutic approaches.

## Methods

### Case Identification

An overview of case identification is presented in Figure 1A. We identified 188 patients whose diagnosis contained the term clear cell ovarian carcinoma (CCOC) from the Edinburgh Ovarian Cancer Database between 1981 and 2016. Following exclusion of 12 patients with recurrent disease or resection following interval debulking surgery (IDS) and 32 with insufficient or no material, pathology review was carried out on 144 cases. The review was performed by two observers (YI, CSH), including a specialist gynecological pathologist (CSH), who applied 2020 WHO criteria [14], leading to exclusion of 42 cases as non-CCOC; these were predominantly tumours that were reported as mixed high grade serous and clear cell carcinomas but on review were considered to represent high-grade serous carcinoma with clear cells. The 102 cases with CCOC morphology were taken forward for IHC to provide further support for the diagnosis; 7 tumours that were WT1 positive were excluded, leaving 95 cases for DNA extraction. Following this, 85 cases were available with sufficient DNA for DNA sequencing. Clinical data, including age, FIGO stage, debulking status, adjuvant chemotherapy, progression-free survival (PFS), and disease-specific survival (DSS), were obtained from the Edinburgh Ovarian Cancer Database; prospectively collected data resource created from tertiary oncology centers in south-east Scotland. Ethical approval was granted by NHS Lothian Research and Development (reference ID 2007/W/ON/29), and the use of human tumour material was approved by the South East Scotland Human Annotated Bioresource (reference 15/ES/0094-SR891).

**Figure 1.**
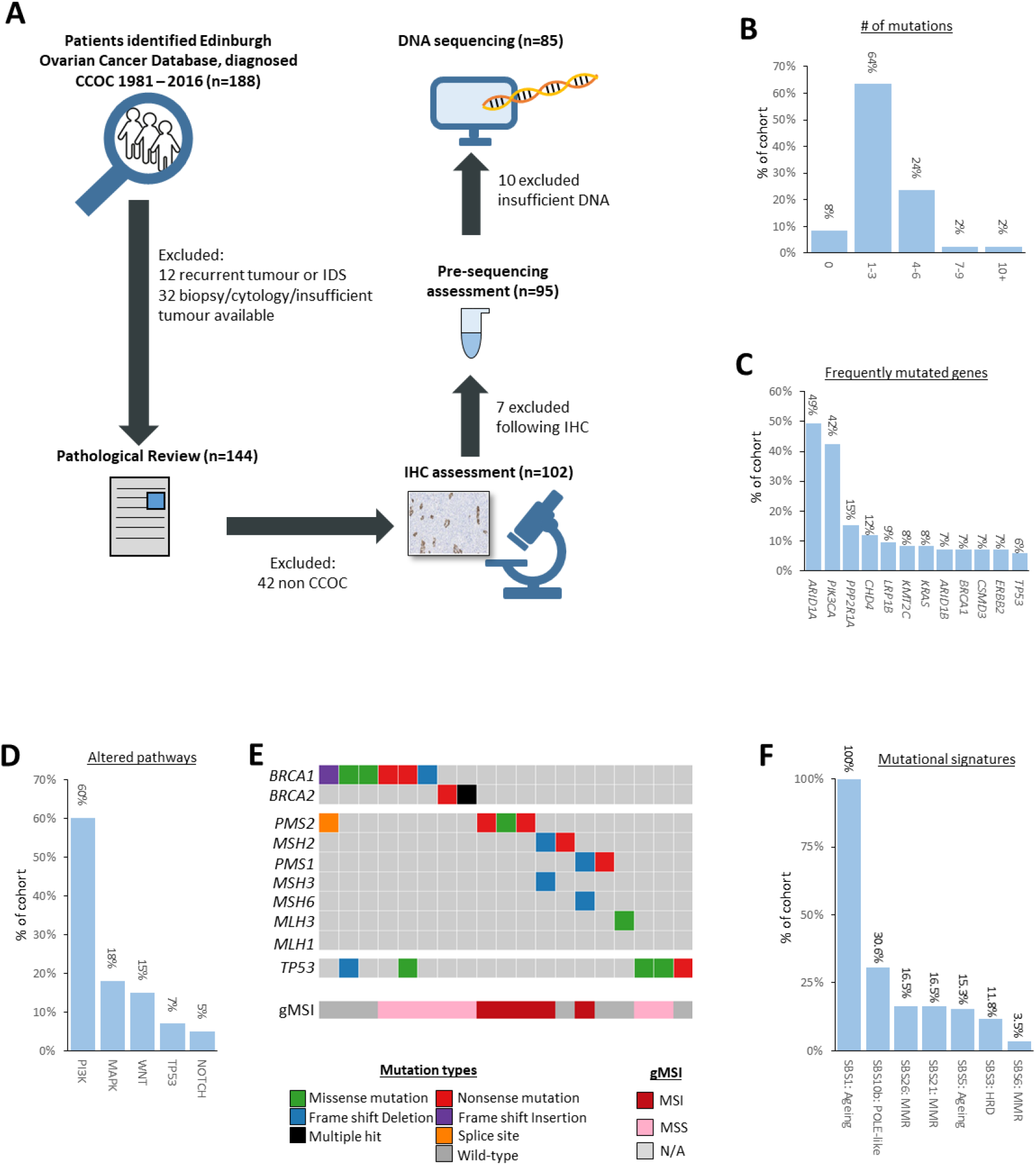
**(A)** Overview of the creation of the clear cell ovarian carcinoma cohort. IDS = Interval Debulking Surgery **(B)** Plot of number of mutations identified per tumour across the panel of sequenced genes, split into bins, as a percentage of the total cohort. **(C)** Plot of mutations detected as a percentage of the total cohort. **(D)** Plot of altered pathways based on mutation data, as a percentage of the cohort. **(E)** Oncoplot of *BRCA1/2*, MMR genes and *TP53* mutations in the cohort. Genomic microsatellite (gMSI) assessment values are shown below. Colour code for mutation types is shown in the legend. **(F)** Plot of the detected mutational signatures, as a percentage of the cohort. SBS = COSMIC Single base substitutions group.

### Immunohistochemistry

IHC was performed on 5μm-thick formalin-fixed, paraffin-embedded (FFPE) tumour sections using antibodies targeting WT1, Napsin A, HNF1β and p53. Immunostaining was automated using the Leica BOND-III autostainer (Leica Biosystems, Newcastle Upon Tyne, UK) and the Bond Polymer Refine Detection kit (Leica Biosystems), following the manufacturer’s protocols. The antibodies and scoring system for IHC are provided in Supplementary Methods.

### Genomic Sequencing

Hematoxylin-Eosin (HE)-stained slides with marked tumour areas were used for microdissection prior to DNA extraction. Sanger sequencing was performed on isolated genomic DNA (gDNA) to screen for TERT promoter mutations. A custom panel-based DNA sequencing assay targeting 68 genes frequently mutated in CCOC, as identified through published whole-exome sequencing studies, was used [2-8]. Sequencing was performed with Unique Molecular Identifiers (UMIs), using the Illumina NextSeq 550 platform (Illumina, San Diego, CA, USA) [Supplementary Table S1]. Targeted sequencing was performed across the cohort with a median per-sample on-target coverage of 355X (range 57X–695X). Data were aligned to the GRCh38 human reference genome using the bcbio 1.0.6 pipeline (see Supporting Methods). Short nucleotide variant (SNV) calling, insertion/deletion (INDEL) prediction, genomic microsatellite instability (gMSI) scoring and pathway analysis were carried out as previously described [15, 16] (see Supplementary methods). Variants were further filtered based on low sequence depth, allele frequency, and non-pathogenic outcomes (see Supplementary methods). Mutational signatures defined by the single base substitution (SBS) signatures in the Catalogue of Somatic Mutations in Cancer (COSMIC: (https://cancer.sanger.ac.uk/cosmic) were detected using the R package “deconstructSigs” [17]. Data analysis and visualization were conducted using the R package maftools [18].

### Immune Cell Infiltration Analysis

Tumour tissue microarrays (TMAs) were constructed using 0.8mm cores from FFPE material derived from genomically characterized CCOC cases. IHC was performed on TMA sections with antibodies for CD3, and CD8 (See supplementary methods for more information). Immunostaining was automated using the Leica BOND III Autostainer. Immunohistochemistry quantification was performed using QuPath v.0.2.0-m8 software [19] with automated scoring validated by two independent human observers (RLH, WG) [Supplementary Figure S1]. For each patient, immune infiltration was calculated as the total number of positive cells across each sample for that patient, divided by the total number of detected cells, yielding a percentage positive cell score. Cases with no evaluable cores were excluded (n=8 for CD3, n=6 for CD8) CD3+ and CD8+ levels were found on a continuous gradient across the cohort. Tumours were therefore classified as “TIL enriched” for CD3+ or CD8+ TILs where the percentage of enriched cells was >5% of the total observable tissue (>85^th^ percentile of data in our cohort), otherwise these were classified as “TIL depleted”.

### Statistical Analysis

All statistical analyses were performed using R version 4.3.2. Categorical variables were compared using Fisher’s exact test (3×2), the Kruskal-Wallis test, or the Cochran-Armitage trend test, as appropriate. Survival analysis was conducted using the Kaplan-Meier method (survival package), with differences between groups assessed by log-rank tests. Univariate Cox proportional hazards models were used to estimate hazard ratios (HR) and 95% confidence intervals (CI), with results visualized using the survminer and forestplot packages. Multivariate Cox regression models were applied to evaluate the independent associations of FIGO stage, age at diagnosis, ARID1A mutational status, and TIL enrichment with progression-free survival (PFS). All p-values were two-tailed, with statistical significance set at p < 0.05.

### Data availability

The genomic data generated for patient tumours presented in this manuscript are stored within University of Edinburgh secure servers and are available upon reasonable request upon review by our internal data access committee.

## Results

### Clinical Characteristics

85 CCOC cases with sufficient DNA for subsequent sequencing were identified from the Edinburgh Ovarian Cancer Database; a prospectively collected data resource created from tertiary oncology centers in the south-east of Scotland [Figure 1A]. All cases were WT1 negative; all were HNF1β positive; and 83 (97.6%) were positive for Napsin A. IHC for p53 showed a wild-type pattern in 81 cases and an aberrant diffuse (mutant) pattern in 4. Clinical characteristics of the CCOC cohort are summarized in Table 1. All patients underwent primary debulking surgery; 71 (87.7% of 81 cases with known debulking status) were debulked to zero macroscopic residual disease (RD). The median follow-up time was 9.62 years (95% CI 9.07-15.45). The median DSS and PFS were 8.52 and 5.44 years.

**Table 1.** Clinical characteristics of clear cell ovarian carcinoma cases.

|  |  | N | % |
| --- | --- | --- | --- |
| Cases |  | 85 | - |
| Age | Median years | 59 | range 30-79 |
| FIGO stage | IA-IC1 | 24 | 28.2 |
|  | IC2-II | 47 | 55.3 |
|  | III-IV | 14 | 16.5 |
| RD following cytoreduction | Macroscopic RD | 10 | 12.3 |
|  | Zero macroscopic RD | 71 | 87.7 |
|  | NA | 4 | - |
| Neoadjuvant chemo | Yes | 0 | 0 |
|  | No | 85 | 100 |
| Adjuvant therapy | Platinum | 26<sup>a</sup> | 30.6 |
|  | Platinum-taxane | 33<sup>b</sup> | 38.8 |
|  | Other platinum combination | 7 | 8.2 |
|  | Other chemotherapy | 5 | 5.9 |
|  | Radiotherapy | 1 | 1.2 |
|  | None (surgery only)<sup>c</sup> | 13 | 15.3 |
| Death details at last follow-up | Alive | 35 | 41.2 |
|  | Deceased, ovarian cancer | 37 | 43.5 |
|  | Deceased, other causes | 13 | 15.3 |
| Diagnosis time period | Pre-2000 | 24 | 28.2 |
|  | 2000-2009 | 25 | 29.4 |
|  | 2010 onwards | 36 | 42.4 |
| Clinical outcome | Median OS | 5.85 | 95% CI 4.33-14.10 |
|  | Median DSS | 8.52 | 95% CI 5.62-NA |
|  | Median PFS | 5.44 | 95% CI 4.19-NA |
| Follow-up time | Median years | 9.62 | 95% CI 9.07-15.45 |

<sup>a</sup>3 cases also treated with radiotherapy; <sup>b</sup>3 cases also treated with radiotherapy; <sup>c</sup>10 stage I cases with complete resection; 2 stage II cases with complete resection; 1 stage III case with rapid deterioration following surgery

### Genomic analysis of the CCOC cohort

Across the entire cohort we report a total of 233 pathogenic variants over 44 of the 69 (64%) evaluated genes, spanning 78 of the 85 samples (92%). The median number of variants per sample was 2, with a range of 0 to 14 [Figure 1B, Supplementary Figure S2 & Supplementary Table S1].

The most frequently mutated genes were *ARID1A* (n=42; 49%) and *PIK3CA* (n=36; 41%), for which 19 cases (22%) had co-occurring mutations across these two genes. Additional mutations were observed in *PPP2R1A* (n=13; 15%), *CHD4* (n=10; 12%), *LRP1B* (n=8; 9%), *KMT2C* (n=7; 8%), and *KRAS* (n=7; 8%) [Figures 1C & 2].

Less common mutations included alterations in *BRCA1, ARID1B, CSMD3*, and *ERBB2* (all n= 6; 7%) [Figure 1C]. Mutations in *TERT* were found in 3 cases (4%), with promoter mutations c.-124C>T and c.-146C>T detected in one and two samples, respectively [Figure 2]. Pathway-level analysis revealed that 51 cases (60%) harbored mutations in the PIK3-AKT-mTOR pathway, followed by perturbations in the MAPK (n=15; 18%), WNT (n=13; 15%), TP53 (n=6; 7%), and NOTCH (n=4; 5%) pathways [Figure 1D].

**Figure 2.**
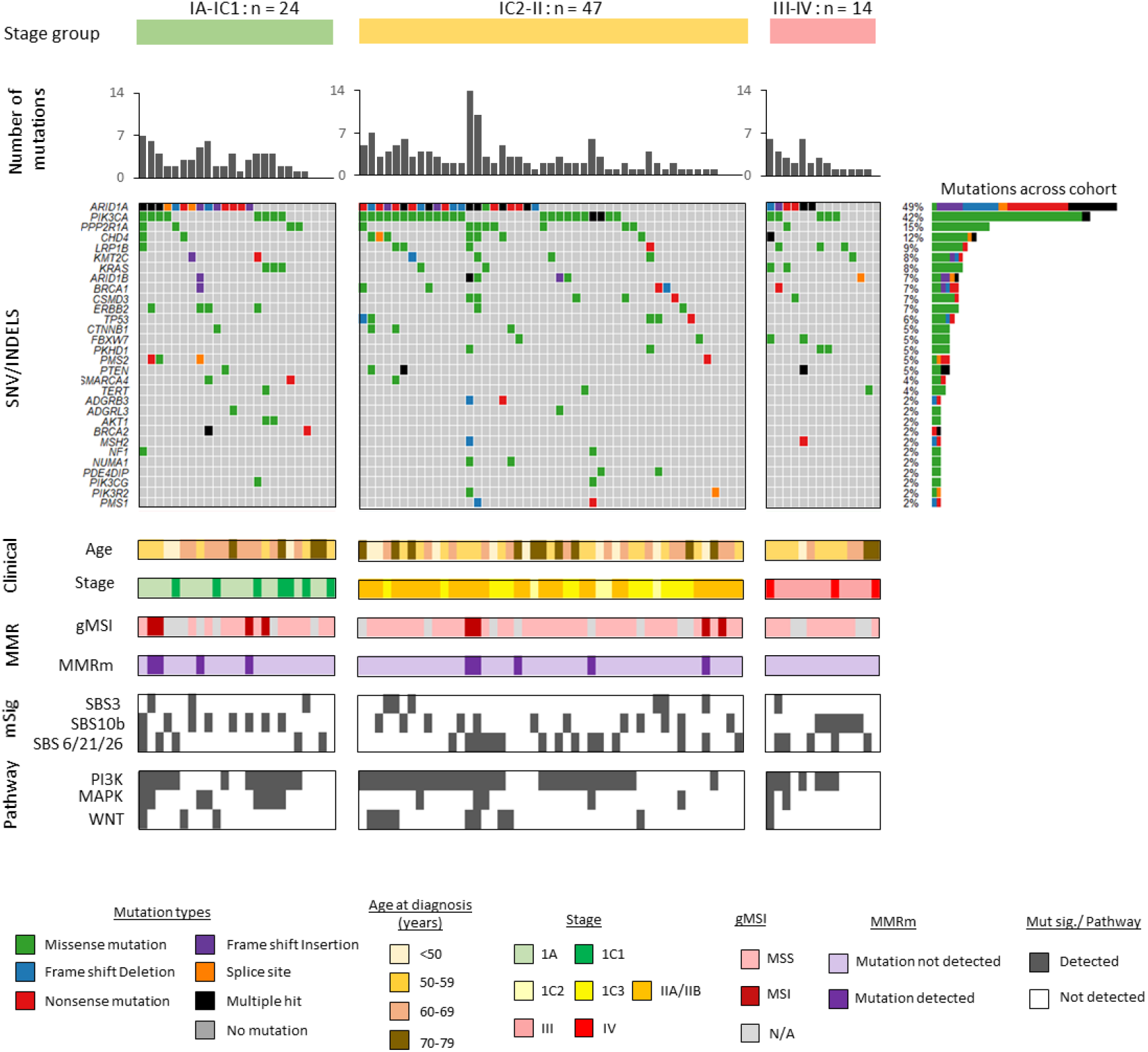
Molecular landscape of the CCOC cohort. Oncoplot of SNV and INDEL events across the CCOC cohort, split by stage group. Upper plot displays number of mutations detected per tumour; SNV = single nucleotide variant, INDEL = short insertions and deletions. Events >2% are displayed with plots of % of detection across cohort shown to the right; Colour defines mutation type. Grey denotes no mutation. Age at diagnosis, FIGO stage, gMSI = genomic micro-satellite instability score (MSS = micro-satellite stable, MSI = micro-satellite instability, N/A = non evaluable), MMR gene mutation, select mutational signatures (“msig”) and pathway perturbation based on mutations are shown below.

Genomic microsatellite instability (gMSI) analysis classified 8 tumours (9%) as microsatellite instability-positive (MSI). Among these, six cases (75%) carried mutations in known mismatch repair (MMR) genes [Supplementary Figure S3]. *BRCA1* and *BRCA2* mutations were detected in 8 cases (combined 9%; *BRCA1* 7%, *BRCA2* 2%) with 2 of the *BRCA1m* cases also containing a mutation in *TP53* [Figure 1E]. These two cases also contained mutations in genes associated with CCOC such as *ARID1A* and *PDE4DIP*. MMR gene mutations were found in 9 cases (11%, “MMRm”; *PMS1, PMS2, MLH3, MSH2, MSH3*, or *MSH6*). Mutations in *BRCA1/2* and MMR genes were mostly mutually exclusive, with co-occurrence observed in only one case (1/16) [Figure 1E].

Mutational signature analysis across the cohort identified an inherent SBS1 signature in all samples, reflecting age-related tumour changes. Additionally, a POLE-like signature (SBS10b) was enriched in 30.6% of cases [Figure 1F]. Three mismatch repair-associated signatures were observed; with SBS26 and SBS21 each found in 16.5% of cases, and SBS6 in 3.5% [Figure 1F].

### Stratification of patients by stage to identify genomic biomarkers

To better understand how the genomic landscape varies by stage in CCOC, we stratified the cohort into three groups based on FIGO stage: IA–IC1 (n=24, 28%), IC2–II (n=47, 55%), and III–IV (n=14, 17%). Stratification revealed no significant differences in mutational load (Kruskal-Wallis non-parametric test), frequencies of gene mutations, frequencies of altered pathways, or presence of mutational signatures (SBS10b, SBS26/21/6, SBS3) (Fisher’s exact tests and Cochran-Armitage test for trends) [Figure 2 & Supplementary Figure S5A]. However, an enrichment in MAPK pathway perturbations was observed in stage IA–IC1 tumours compared to later stage tumours (Fisher’s exact test, *P* = 0.05 [Figure 2A and Supplementary Figure S4]. A trend was noted in the reduction of the proportion of MSI-positive cases by gMSI as stage increased with 4/17 (24%) stage IA–IC1, 4/39 (10%) stage IC2–II, and 0/10 (10%) of stage III– IV evaluable tumours; Cochran-Armitage test for trend *P* = 0.059) [Supplementary Figures S4B & S4C]. Although not reaching significance, there is an enrichment of mutations within the ARID domain and receptor-dependent activation domain of ARID1A in low-stage tumours (FIGO stages IA–IC1) compared to advanced-stage tumours (FIGO stages IC2–II and III–IV). Mutations in these domains were found in 6/24 (25%) cases in stages IA–IC1, 7/47 (14.9%) cases in stages IC2–II, and 0/14 (0%) cases in stages III–IV (Fisher’s exact test *P* = 0.237, Cochran-Armitage test for trend *P* = 0.076 [Supplementary Figures S5B & 5C].

### Tumour infiltrating immune cells in CCOC

We next conducted a quantified assessment of per-patient infiltration burden through analysis of tumour tissue microarrays (TMAs) using IHC for each tumour sample. The degree of CD3+ and CD8+ tumour-infiltrating lymphocytes (TILs) was highly heterogeneous across CCOC cases, with a median percentage of CD3+ cells at 0.28% (interquartile range [IQR] 0.09–2.04%) and CD8+ cells at 0.18% (IQR 0.03–1.52%) [Figure 3A and Supplementary Figure 6].

**Figure 3.**
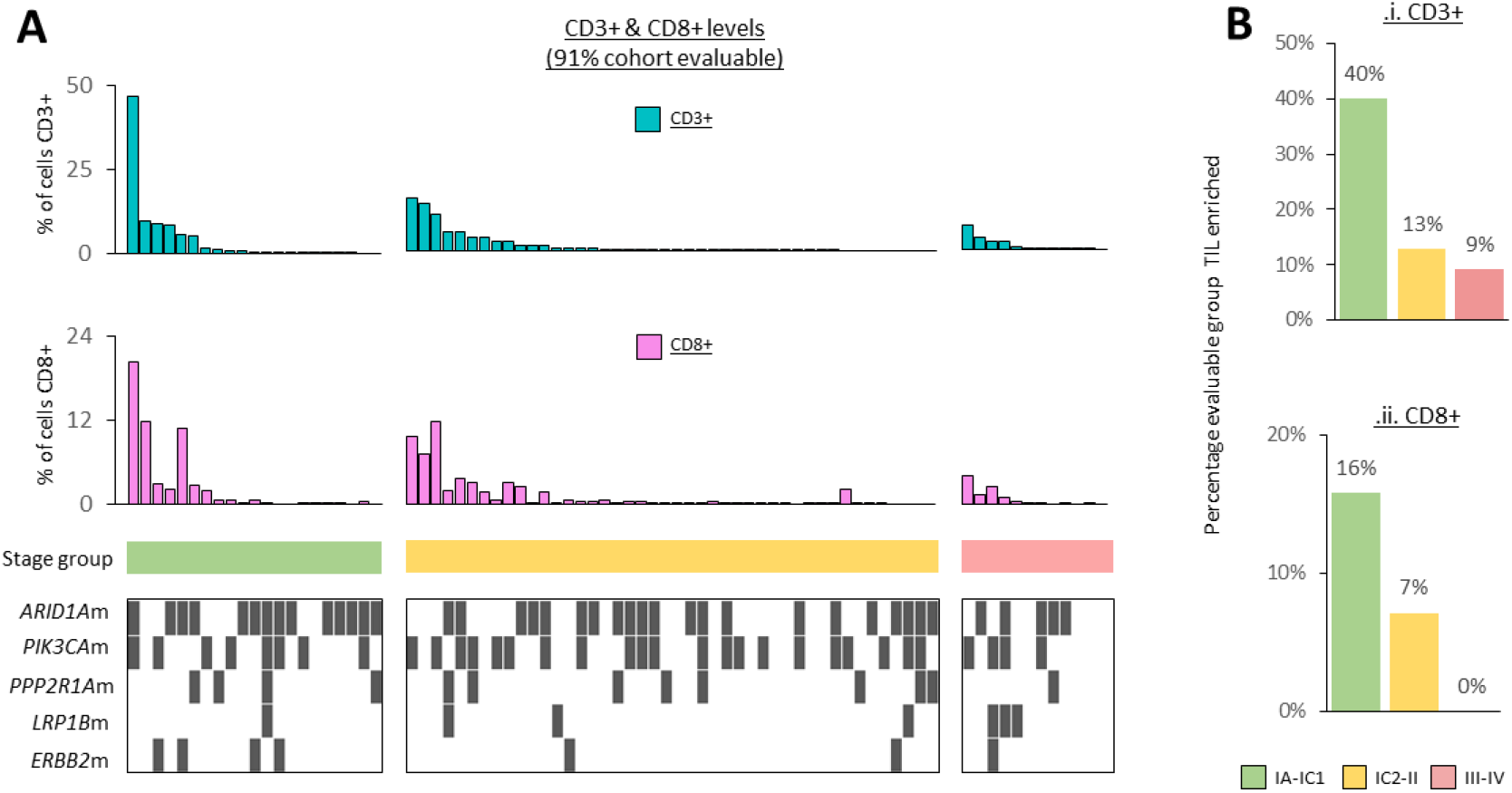
Immune cell infiltration markers CD3 and CD8 across the cohort. (**A)** Bar plot of percentage of CD3 (light blue) and CD8 (pink) detected in a given tumour section, ranked from high to low based on CD3+ levels, split by the stage groups. Only tumours with evaluable CD3 and CD8 data are shown, representative of 91% of the cohort. Select genomic mutational events are plotted below. **(B)** Plot of the percentage of tumours defined as “enriched” in (i) CD3 or (ii) CD8 (≥5% overall cell count) across evaluable samples, split by stage group.

No significant differences in CD3+ or CD8+ cell infiltration levels were observed when stratified by the mutational status of frequently mutated genes, nor when split across the three stage groups (Fisher’s exact tests and Cochran-Armitage test for trends); although notable trends were observed demonstrating a potential enrichment for both CD3+ and CD8+ve enriched cases in evaluable IA–IC1 tumours (40% & 16% respectively) compared to later stage tumours (CD3+: IC2–II = 13%, III–IV = 9%, CD8+: IC2–II = 7%, III–IV = 0%) [Figure 3B & Supplementary Figure 7].

### Prognostic Risk Factors for Low and Advanced-Stage Clear Cell Ovarian Carcinoma

Univariate analysis of progression-free survival (PFS) across the cohort, stratified by tumour stage groups, revealed a significant association between stage at diagnosis and PFS, as expected. Using IA–IC1 as the reference group, patients with IC2–II stage tumours demonstrated a hazard ratio (HR) for reduced PFS of 3.5 (*P* = 0.023), while those with stage III–IV tumours demonstrated an HR of 90.9 (*P* < 0.001) [Supplementary Figure S8]. Univariate analysis of *ARID1A* mutation status did not reveal a significant relationship with PFS across the whole cohort (*ARID1A*m HR = 0.73, *P* = 0.326; 95% CI 0.38-1.4) [Supplementary Figure S9]. Despite trends in both, low levels of CD3+ve TILs, but not CD8+ve TILs, were significantly associated with reduced PFS across the entire cohort (CD3+ depleted HR = 4.4, *P* = 0.042, 95% CI 1.1-18; CD8+ depleted HR = 4.6, 95% CI 0.63-34, log-rank *P* = 0.098) [Supplementary Figure S10]. Multivariable analysis of PFS across the whole cohort incorporating tumour stage by grouping IA–IC1 as “low stage” and IC2–IV as “advanced stage” alongside age at diagnosis (years), *ARID1A* mutation status, CD3+ TIL depletion and CD8+ TIL depletion, revealed no significant molecular marker associated with relapse across the cohort as a whole [Supplementary Figure S11].

In order to identify risk factors associated with earlier-stage disease, which is particularly challenging to manage clinically, we retained the stratified groups of “low stage” (IA-IC1) and “advanced stage” (IC2-IV) and investigated molecular markers across these subgroups. We note worse PFS in *ARID1A*wt cases, exclusively in IA–IC1 stage tumours (*ARID1A*wt; HR = 7.2, P = 0.088; 95% CI 0.74-72) [Supplementary Figure S12]. Combining CD3+ TIL depletion and *ARID1A*wt status as a composite risk factor improves upon this finding with increased recurrence risk specifically in stage IA–IC1 tumours (HR = 11.7, *P* = 0.05; 95% CI 1-138) [Figure 4]. No such association was observed in more advanced stages (HR = 1.1, *P* = 0.75; 95% CI 0.55-2.3) or when investigated across the whole cohort unstratified by stage (HR = 1.8, *P* = 0.98; 95% CI 0.9-3.5) [Figure 4]. Although we note that the percentage of patients receiving adjuvant therapy was greater in late stage CCOC (58.3% low stage, 91.8% of late stage), there was no significant difference when the cohort was stratified by combined risk factor presence (Chi-squared *P* = 0.449).

**Figure 4.**
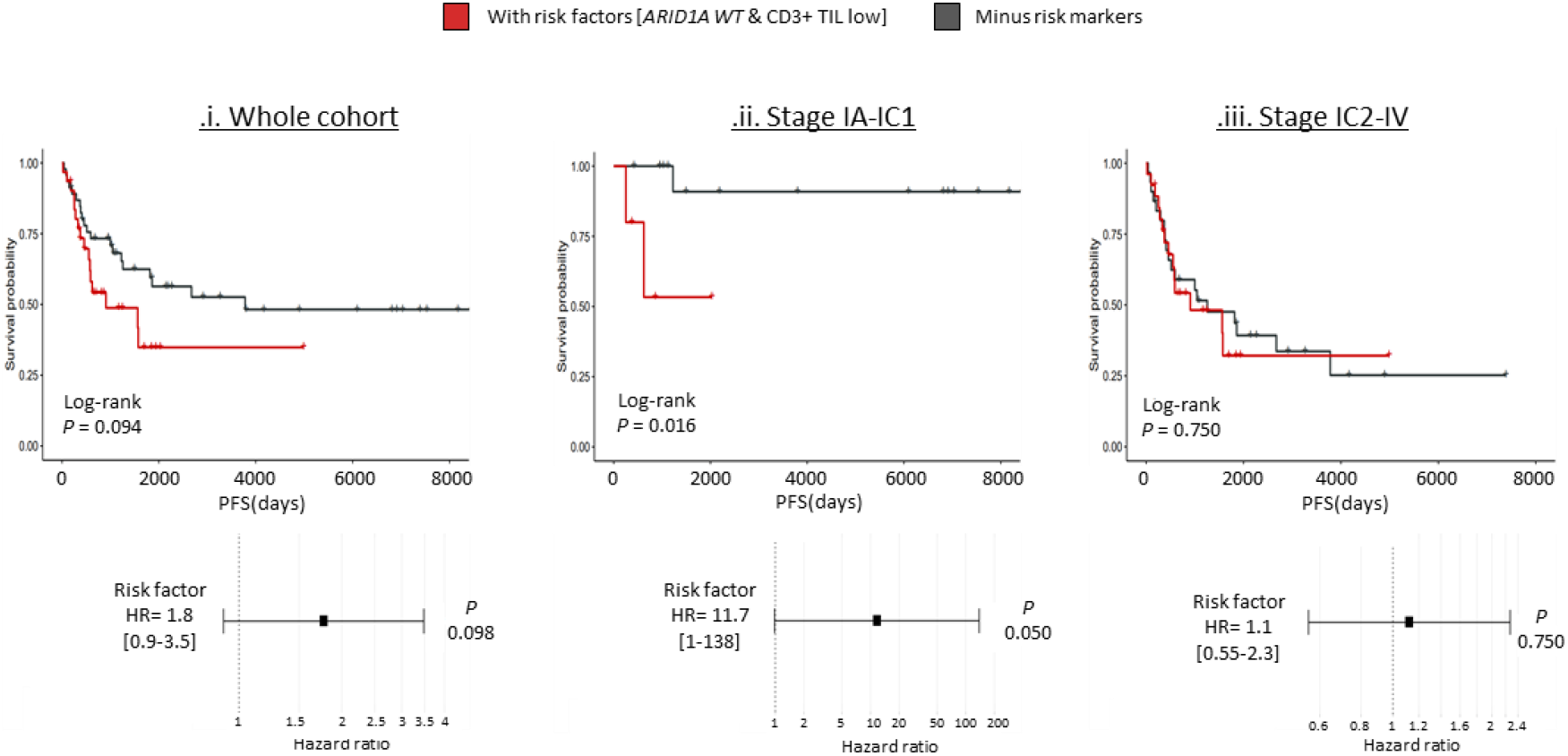
Identification of markers associated with poor outcome in low stage CCOC. Kaplan-Meier survival analysis and hazard ratio plot for progression-free survival (PFS) in the study cohort when stratified by combined risk factors for (i) the whole cohort, (ii) low-stage IA-IC1 or (iii) advanced stages (IC2-IV). (Upper plots) Kaplan-Meier curve comparing PFS between risk factor groups, with statistical significance assessed by log-rank test. (Lower) Forest plot displaying the hazard ratio (HR) from a univariate Cox proportional hazards model for groups, with 95% confidence intervals.

As such, A combined lack of *ARID1A* mutation with low CD3+ve tumour infiltration levels show promise as a method to identify a subgroup of low-stage CCOC patients at increased risk of recurrence.

## Discussion

Here we provide a detailed examination of genomic and immune biomarkers associated with relapse risk in a well-defined cohort of CCOC, with rich associated clinical annotation, with a particular emphasis on low-stage disease (FIGO stage IA–IC1). Our results highlight the prognostic importance of *ARID1A* mutational status and tumour-infiltrating lymphocyte (TIL: CD3+ and CD8+ T cells) levels. Through integrating these genomic and immune features, we identify a subgroup of patients with *ARID1A* wild-type (WT) tumours and low CD3+ve TIL levels who are at a significantly elevated risk of recurrence. These results offer important insights into potential avenues for patient stratification and therapeutic intervention, underscoring the need for tailored approaches in low-stage CCOC.

This study builds on recent efforts to characterize the genomic landscape of CCOC [2-5, 8]. In agreement with these studies our analysis reveals a heterogeneous mutational landscape, with frequent alterations in genes such as *ARID1A* (49%), *PIK3CA* (41%) and less frequent mutation in *PPP2R1A* (15%), and *CHD4* (12%) as well as a wider perturbation across the PI3K-AKT-mTOR (60%) and MAPK (18%) signaling pathways. Genomic microsatellite instability (gMSI) and mismatch repair (MMR) deficiency were detected in a small proportion of cases (9%), highlighting potential avenues for therapeutic exploration. Mutational signature analysis identified contributions from processes associated with defective homologous recombination repair and polymerase epsilon (POLE) mutations in select cases, further emphasizing the molecular diversity of CCOC.

Our findings on *BRCA1/2* and MMR mutations also align with previous reports for CCOC, yet also provide additional insight into their patterns of mutual exclusivity [2-4]. We report a low prevalence of both *BRCA1/2* mutations (9% in our study) and MMR gene mutations (11% in our study) which were largely mutually exclusive events, suggesting potential distinct molecular pathways with BRCA-associated and MMR-deficient tumours representing divergent subgroups within CCOC. In agreement with previous reports [8], we note that a small proportion of our cohort (9%) are associated with MSI, typically in cases with an observed mutations in a MMR gene (75% of gMSI cases). We also observed a low prevalence of *TP53* mutations (6%), consistent with levels noted in previous reports (7-16%) [3, 4, 8], reinforcing the rarity of *TP53* alterations in cohorts of CCOC that are robustly reviewed to exclude occult high grade serous carcinomas demonstrating clear cell change. We identified two cases with concurrent *BRCA1* and *TP53* mutations. These two cases also contained mutations in genes associated with CCOC such as *ARID1A* and *PDE4DIP*. Given the rigorous histotype confirmation of our cohort through WT1 negativity and Napsin A/HNF1β positivity, we are confident that these mutation patterns reflect true CCOC biology rather than misclassification with other ovarian cancer subtypes.

Interestingly, we note an enrichment of MAPK pathway mutations in low-stage tumours, present in 33% of IA–IC1 cases compared to 14% in advanced-stage tumours. This suggests a potential interaction between MAPK pathway alterations and *ARID1A* mutations that may influence the tumour microenvironment. The MAPK pathway is known to regulate immune cell recruitment and inflammatory responses [20, 21]. Targeted therapies focusing on the PIK3-AKT-mTOR and MAPK pathways may emerge as additional strategies, given the high prevalence of these alterations in the study cohort (60% and 18%, respectively).

Our study also highlights the heterogeneity of immune infiltration in CCOC. We found that TIL enrichment was more common in low-stage (IA–IC1) tumors (29%) than in advanced stages (18% in IC2–II and 17% in III–IV), suggesting a more active immune response in low stage disease. Additionally, whilst recent reports have found no correlation between immune infiltration in treatment-naïve tumors and overall survival [12], our analysis suggests that immune infiltration may have prognostic relevance, particularly in low-stage CCOC.

A recent study using multiplex immunofluorescence analysis reported that T-cell infiltration was significantly higher in adjacent nonneoplastic regions than in tumor tissue, with a greater differential in CD8+ cell density between these regions being associated with poorer PFS (p = 0.042) [22]. This aligns with our observation that lower TIL levels in low-stage CCOC are linked to an increased risk of recurrence, suggesting that immune exclusion mechanisms may play a role in tumor progression. Furthermore, this study found that high CD8+ T-cell expression in adjacent stromal and nonneoplastic regions correlated with worse PFS, reinforcing the notion that immune cell localization as well as absolute abundance, is a key determinant of clinical outcomes [22].

It has also been reported that PD-L1 and T cell expression increases at recurrence in CCOC, with no significant differences in these markers between platinum-sensitive and platinum-resistant disease, although higher TIL levels were observed in platinum-sensitive cases [12]. Whilst we did not directly assess platinum sensitivity, our finding that TIL-depleted tumors have worse PFS supports the idea that immune-infiltrated tumors may have better outcomes and potentially better treatment responses. Together, these results suggest that immune contexture in CCOC evolves over time, with potential implications for recurrence risk and treatment strategies.

The findings from the PEACOCC trial, which demonstrated clinical benefit with pembrolizumab in advanced clear cell gynecological cancers, highlighted the potential for PD-1 inhibition, particularly in the setting of mismatch repair (MMR)-proficient tumors, where pembrolizumab showed a modest progression-free survival (PFS) benefit [23]. This study agrees with our findings suggesting that immune evasion mechanisms, such as low TIL levels, could influence treatment responses and recurrence risk. These findings underscore the importance of immune profiling in CCOC and suggest that immune modulation strategies, such as PD-1 inhibition, could hold promise in specific subgroups of CCOC patients.

Our findings also agree in part with recent large-scale analyses of ARID1A and immune infiltration proteins in CCOC [24]. The OTTA/COEUR study, which included 545 CCOC cases, found no significant association between ARID1A loss and CD8+ TIL status, both by IHC. This study also reported no notable prognostic impact of ARID1A protein loss, across their entire CCOC cohort, which agrees with our findings prior to stratification by stage. This study also reported no notable prognostic impact of CD8+ TIL status across CCOC samples.

We report that combining the TIL and genomic results allow for the identification of a high-risk subset of low-stage tumours characterized by low TIL levels and *ARID1A* WT status, which exhibited a markedly higher risk of recurrence (HR = 8.9, P = 0.059). This finding supports the hypothesis that immune evasion mechanisms may be particularly prominent in *ARID1A* WT tumours, which are less likely to exhibit the increased neoantigen burden and DNA damage repair defects associated with *ARID1A* mutations [25]. Our findings also align with emerging evidence on the role of *ARID1A* in modulating immune responses, where *ARID1A*-deficient OCCC cells were seen to exhibit greater sensitivity to TCR-T cell therapy, supporting the notion that *ARID1A* mutations may create a more immunogenic tumor microenvironment [26]. This complements our observation that *ARID1A*wt tumors in low-stage disease may be less immunologically active, contributing to worse outcomes. Together, these findings highlight the potential of immune-based therapies, particularly for *ARID1A*-deficient CCOC, and reinforce the need to consider *ARID1A* status when evaluating immunotherapeutic strategies.

A significant strength of this study is the use of a well-characterized cohort with detailed clinical and molecular data, enabling robust stratification by stage and comprehensive integration of genomic and immune analyses. However, the study’s relatively small sample size, particularly in subgroup analyses, limits the statistical power to detect subtle associations. For example, we were unable to use the CD8+ marker as a composite risk factor in low stage analysis due to the low number of cases demonstrating CD8+ enrichment and relapsed state when stratified by stage. Future research should validate these findings in larger, prospective cohorts and explore the interactions between *ARID1A* mutations, MAPK pathway alterations, and immune infiltration. Longitudinal studies tracking dynamic changes in immune infiltration during disease progression could provide additional insights into the role of the tumour microenvironment in relapse risk.

## Conclusion

This study identifies *ARID1A* WT status and CD3+ TIL depletion as prognostic factors for relapse in low-stage CCOC. By integrating genomic and immune biomarkers, the findings lay a foundation for improved risk stratification and the development of personalized therapeutic strategies. The study underscores the potential of combining genomic and immune analyses to guide clinical decision-making and improve outcomes in this challenging ovarian cancer subtype.

## Acknowledgments

We are grateful to the NHS Lothian Department of Pathology, Edinburgh Experimental Cancer Medicine Centre, the Edinburgh Ovarian Cancer Database and the NRS Lothian Human Annotated Bioresource for their ongoing support. The DNA sequencing described here was supported by the Edinburgh Clinical Research Facility, Western General Hospital, Edinburgh, UK. We thank Alison Meynert and Colin Semple (MRC Human Genetics Unit, Institute of Genetics and Cancer, University of Edinburgh) for general support with informatics pipelines.

## Author contributions

YI: data curation, investigation, methodology, manuscript writing – review and editing. RH: data curation, investigation, methodology, manuscript writing – review and editing. MC: data curation, project administration, manuscript writing – review and editing. ST: manuscript writing – review and editing. CB: Sample collection. IC: methodology. WG: investigation, manuscript writing – review and editing. TR: data curation. RN: TBC. CSH: conceptualization, investigation, methodology, supervision, writing – original draft, review and editing. CG: conceptualization, supervision, methodology, writing – review and editing. AO: conceptualization, supervision, methodology, writing – review and editing. JPT : conceptualization, investigation, methodology, sequencing data processing, bioinformatics analysis, data management, manuscript writing – original draft, review and editing.

