## Supplementary Information for "Combined genomic and molecular analysis defines prognostic markers of relapse in stage IA-IC1 clear cell ovarian carcinoma"

#### Contents

Page 2. A. Supporting Methods

Page 7. B. Supporting Figures

Page 19. C. Supporting Table

### A. SUPPORTING METHODS

##### 1. Immunohistochemistry

IHC was performed on 5 µm-thick formalin-fixed, paraffin-embedded (FFPE) tumour sections, automated using the Leica BOND-III autostainer (Leica Biosystems, Newcastle Upon Tyne, UK) and the Bond Polymer Refine Detection kit (Leica Biosystems), following the manufacturer's protocols. The following antibodies and scoring systems were used: WT1 (Dako), clone 6F-H2 at primary dilution of 1:1000 with positive nuclear staining categorized as present in >1% of tumour cells; p53 (Dako), clone DO-7 at primary dilution of 1:50, interpreted as follows: the aberrant diffuse pattern, categorized as diffuse staining in ≥ 70% of tumor cells; the aberrant null pattern if tumour cells were negative with positive internal control stromal cells; and the wild type pattern where staining was variable in intensity between tumour cells; Napsin A (Leica), clone IP64 at a predilute primary dilution with cytoplasmic staining categorized as positive if present in >1% of tumour cells and negative if below this; HNF1B (Atlas), clone HPA002083 at primary dilution of 1:200 with nuclear staining categorized as positive if present in >1% of tumour cells and negative if below this.

##### 2. DNA extraction

Macrodissection was performed using up to six 10µm FFPE sections. Hematoxylin-Eosin (HE) stained slides, in which tumour areas with cellularity at least 30% or more had been marked by an expert gynecological pathologist (CSH), were used as a guide for macrodissection. DNA extraction was performed using the QIAmp DNA FFPE Tissue Kit (QIAGEN, Crawley, UK) and Deparaffinization Solution (QIAGEN) following the manufacturer's instructions. The quality and quantity of the DNA were assessed spectrophotometrically using DS-11 FX (Denovix, Wilmington, Delaware, USA). The quantity of the DNA was further assessed fluorescently using by Qubit ds DNA BR Assay kit (Thermo Fisher Scientific, Waltham, MA, USA) and DS-11 FX (Denovix).

##### 3. Sanger sequencing for TERT promoter

Sanger sequencing was performed to screen for *TERT* promoter mutations. The *TERT* promoter including the two mutation hotspots (-124 and -146 bp upstream from the ATG start site) was amplified by polymerase chain reaction (PCR) with the pair of primers: 5'-CAGCGCTGCCTGAAACTC-3' (Forward) and 5'-GTCCTGCCCCTTCACCTT-3' (Reverse). PCR was performed using Multiplex PCR Kit (QIAGEN). PCR QC via gel electrophoresis products were purified by ExoSAP IT (Thermo Fisher Scientific). The sequencing reaction was performed using BigDye Terminator (Applied Biosystems, Foster City, CA, USA). The protocol of the sequencing reaction was as follows; 96°C for 5 minutes; 40 cycles at 96°C for 30 seconds, 50°C for 45 seconds, 60°C for 4 minutes. The products of the sequencing reactions were purified with DyeEx 2.0 Spin Kit (QIAGEN) following the manufacturer's instructions and were sequenced with the 3130xl Genetic Analyzer (Applied Biosystems) or 3730 DNA Analyzer (Applied Biosystems). Mutational analysis was performed using Mutation Surveyor V5.0.1 (SoftGenetics LLC, PA, USA).

**4. Panel-based DNA sequencing**

For library preparation, 200ng of each DNA sample was sheared to 300-bp fragments with the Covaris E220 Evolution (Covaris, Woburn, MA, USA). Sheared DNA underwent end-repair, A-tailing, and ligation with multiple indexing adaptor (Integrated DNA Technology, San Jose, CA, USA) using the KAPA Hyper Prep Library kit (Kapa Biosystems, Wilmington, MA, USA). Adaptor-ligated DNA were amplified by 8 cycles of PCR using KAPA HiFi HotStart Ready Mix (Kapa Biosystems) and KAPA Library amplification Primer Mix (Kapa Biosystems) and purified using KAPA Pure Beads (Kapa Biosystems). 100ng of up to 16 libraries with unique indexes were pooled together and subjected to target enrichment performed using xGen Hybridization and Wash Kit (Integrated DNA Technology) containing Human cot-1 DNA, and Dynabeads™ M-270 streptavidin. Human cot-1 DNA and xGen Universal Blocking Oligos (Integrated DNA Technology) complementary to our custom adaptor were used to block nonspecific binding of library DNA to targeted probs. Targeted regions were hybridized with customized xGen lockdown probes (Integrated DNA Technology) designed to capture 68 genes (Supporting Table S1) recurrently mutated in CCOC and then captured with Dynabeads™ M-270 streptavidin. Captured libraries were amplified by 13 cycles of PCR using KAPA HiFi Hotstart ReadyMix and xGen Library Amplification Primer (Integrated DNA Technology) and purified using Agencourt AMPure XP beads (Beckman Coulter, Brea, CA, USA). Final libraries were quantified using the Qubit DNA HS assay kit (Invitrogen, ThermoFisher Scientific, Paisley, U.K.), and the size distribution of libraries was assessed using Agilent 2100 Bionanalyser (Agilent Technologies, Stockport, UK). Sequencing was performed using the Illumina NextSeq 550 platform (Illumina, San Diego, CA, USA) generating 150-bp paired-end reads.

**5. Mapping of sequenced reads**

Base calling and quality were assessed using FASTQC. Data were processed with the bcbio-nextgen python toolkit for fully automated high throughput sequencing analysis (see https://github.com/bcbio/bcbio-nextgen for full documentation and informatic pipelines). Raw sequence data was mapped to the hg38 genome build using the Burrows–Wheeler alignment algorithm 0.7.17 with unique molecular identifier (UMI) sequences collapsed as part of the BCBIO pipeline (median UMI duplication reduction factor of 3.37X).

**6. Informatic processing, variant calling and genomic assessment**

As part of the BCBIO pipeline, variant calling was carried out on mapped, UMI compressed BAM files using a majority vote approach from three variant caller algorithms; VarDict [1], Mutect2 [2], Freebayes [3]. Filtering for FFPE and oxidation artifacts was applied using the GenomeAnalysisToolkit (GATK) (CollectSequencingArtifactMetrics and FilterByOrientationBias). Resulting variant call (VCF) files were analysed in R using the maftools package [4]. Datasets were filtered to remove common population variants using the 1000 Genomes reference datasets (1000 genomes phase 1 SNP and InDel dataset; http://www.internationalgenome.org/) and the Exome Aggregation consortium (ExAC) reference datasets (ExAC.0.3.GRCh38 : http://exac.broadinstitute.org/).

Variants of known function were flagged using the NCBI ClinVar database ; variants of unlikely functional significance were filtered using the Polymorphism Phenotyping (PolyPhen) and Sorting Intolerant from Tolerant (SIFT) prediction tools using a read out of one or both tools. Variants with <10% variant allele frequency or a read coverage of <20X were removed.

Data analysis and visualizations were carried out using the R package Maftools [4]. Common SNVs were analysed with the plot function *Oncoplot*. Visual representations of mutational position and associated amino acid changes within genes of interest were drawn using the *lollipopPlot* function. Mutual exclusivity and co-mutations were assessed with the *somaticInteractions* function.

Assessment of genomic microsatellite instability (gMSI) was performed using MSIsensor2 (https://github.com/niu-lab/msisensor2). For some samples, no analysis could be conducted due to insufficient data, likely owing to the panel-based nature of the sequencing, and these samples were labeled as "N/A". Mutational load was defined as the number of mutations present following the aforementioned filtering steps. Mutational signature analysis was carried out using the deconstructSigs R package [5], which has previously been employed to investigate mutational signatures across a range of cancers including ovarian tumours [6]. In brief, stage A filtered mutation data was decomposed into a matrix in the trinucleotide context (matrix of n rows x 96 single base substitution (SBS) class columns, n being number of sequenced samples). The "mut.to.sigs.input" function was used to construct the appropriate input data structure. Then we used "whichSignatures" to determine the COSMIC signatures (v3.3 - June 2022) present in the tumour samples.

**7. Immune cell infiltration analysis**

Tumour tissue microarrays (TMAs) were constructed using 0.8mm cores of FFPE material from genomically characterized CCOC cases. Coring was guided by an H&E stained slide marked by an expert pathologist (CSH) to identify regions of high tumour cellularity. Four cores per patient were used to construct the TMA. IHC was performed on 4µm TMA sections mounted on positively charged slides, using the Leica BOND III Autostainer with BOND ready-to-use anti-CD3 and BOND ready-to-use anti-CD8. Human tonsil was used as a positive control for CD3 and CD8. IHC was quantified using QuPath v.0.2.0-m8 [7] . For each core, tumour area was marked as a region of interest and positive cell infiltration was quantified using the whole cell positive cell detection protocol. Automated scoring was validated using two human observers (RLH,WG); strong correlation was observed between all observers (human observer 1 vs human observer 2 vs machine; Spearman's rho>0.85, P<0.0001 for all comparisons) (Supplementary Figure S1). For each patient, immune infiltration was calculated as the total number of positive cells across each sample for that patient, divided by the total number of detected cells, yielding a percentage positive cell score. Tumours were classified as "TIL enriched" for CD3+ or CD8+ TILs where the percentage of positive cells was >5% (>85th percentile), otherwise these were classified as "TIL depleted".

#### B. SUPPORTING FIGURES

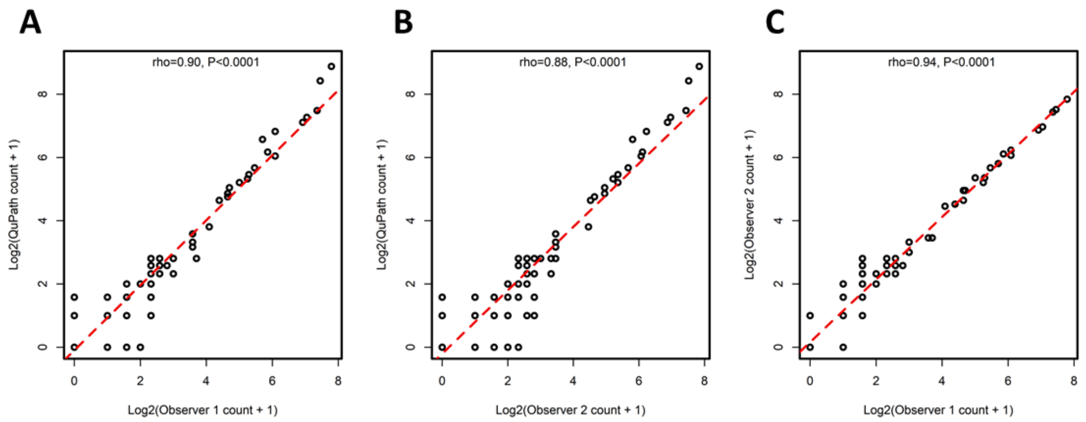


**Figure S1.** Correlation of scoring between (A) Observer 1 and QuPath , (B) Observer 2 and QuPath and (C) Observer 1 and Observer 2.

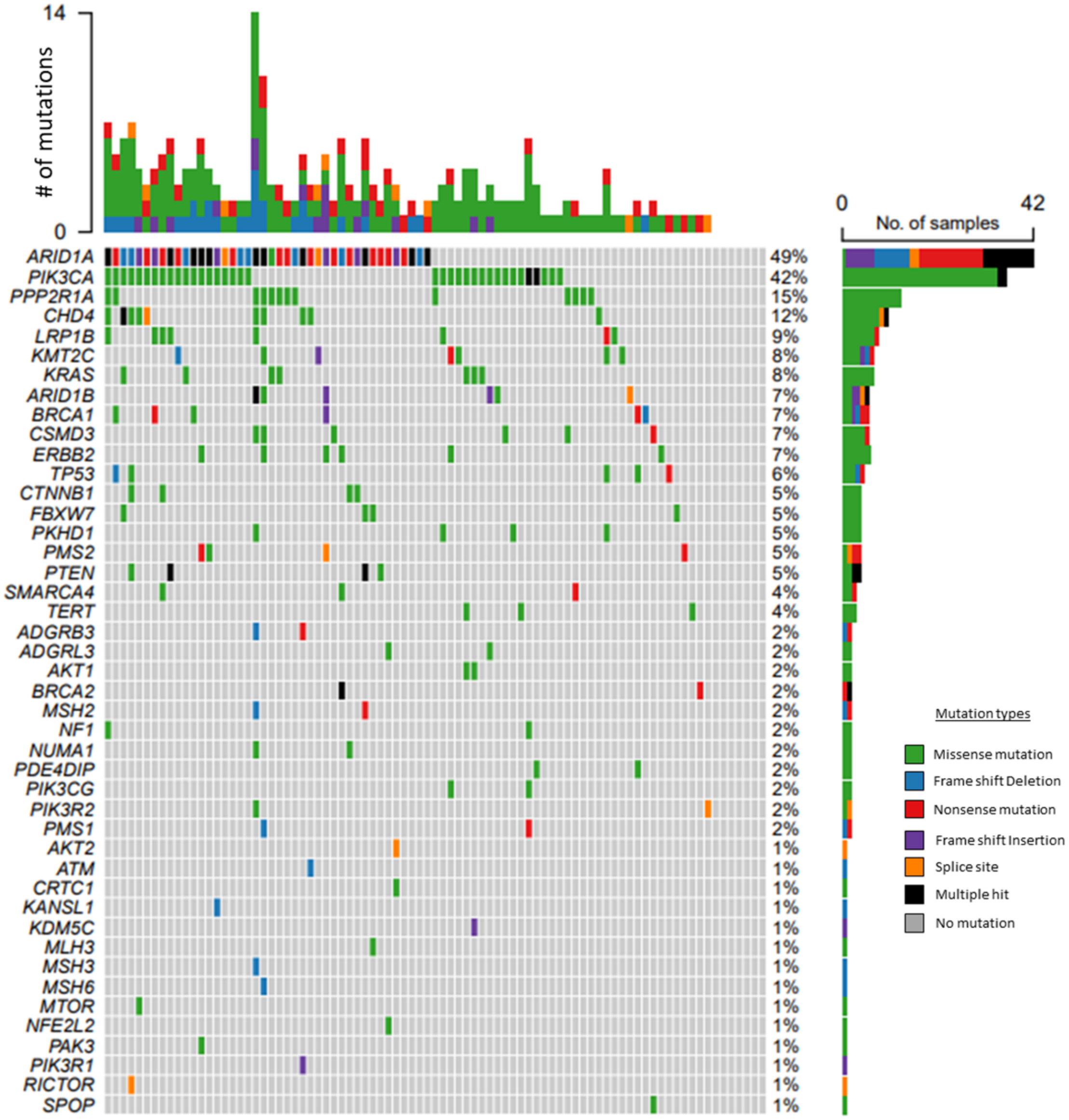


**Figure S2.** Oncoplot for all of the mutated genes detected through SNV analysis. Colour code defines mutation type. Grey denotes no mutation. Total mutation count and type for each tumour are shown above.

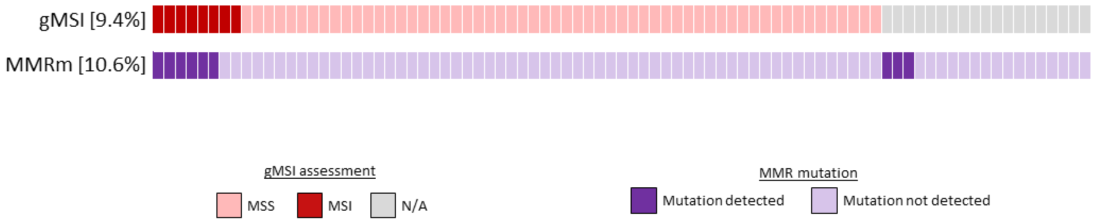


**Figure S3.** Genomic microsatellite (gMSI) assessment across the cohort vs MMR mutation. MMRm genes included *MLH1, MLH3, MSH2,MSH3,MSH6, PMS1* & *PMS2.*

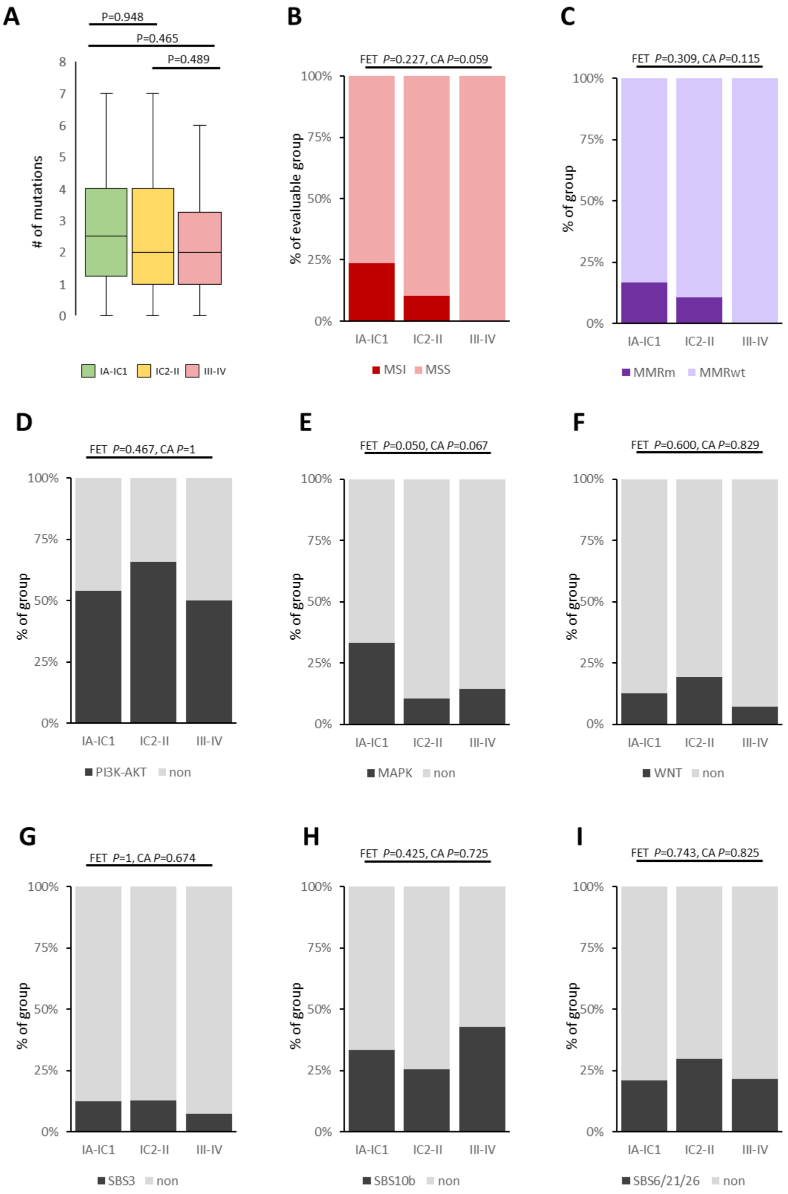


**Figure S4.** Plots of genomic features split by FIGO classes. (A) Boxplot of number of filtered mutations detected across the panel, split by stage group. (B) Stacked bar plot of tumours classified as microsatellite instability positive (MSI) or stable (MSS) by genomic instability assessment. (C) Stacked bar plot of tumours classified as containing at least one mutant mismatch repair gene (MMRm). (D-F) Stacked bar plot of altered pathways for PIK3-AKT (D), MAPK (E) and WNT (F). (G-I) Stacked bar plot of detectable mutational signatures for SBS3 (G), SBS10b (H) and combined SBS6/21/26 (I). P-values are shown for all plots; Kruskal-Wallis non-parametric test for boxplot, 3x2 Fishers Exact Test ("FET") and Cochran-Armitage test for trend ("CA") for stacked plots.

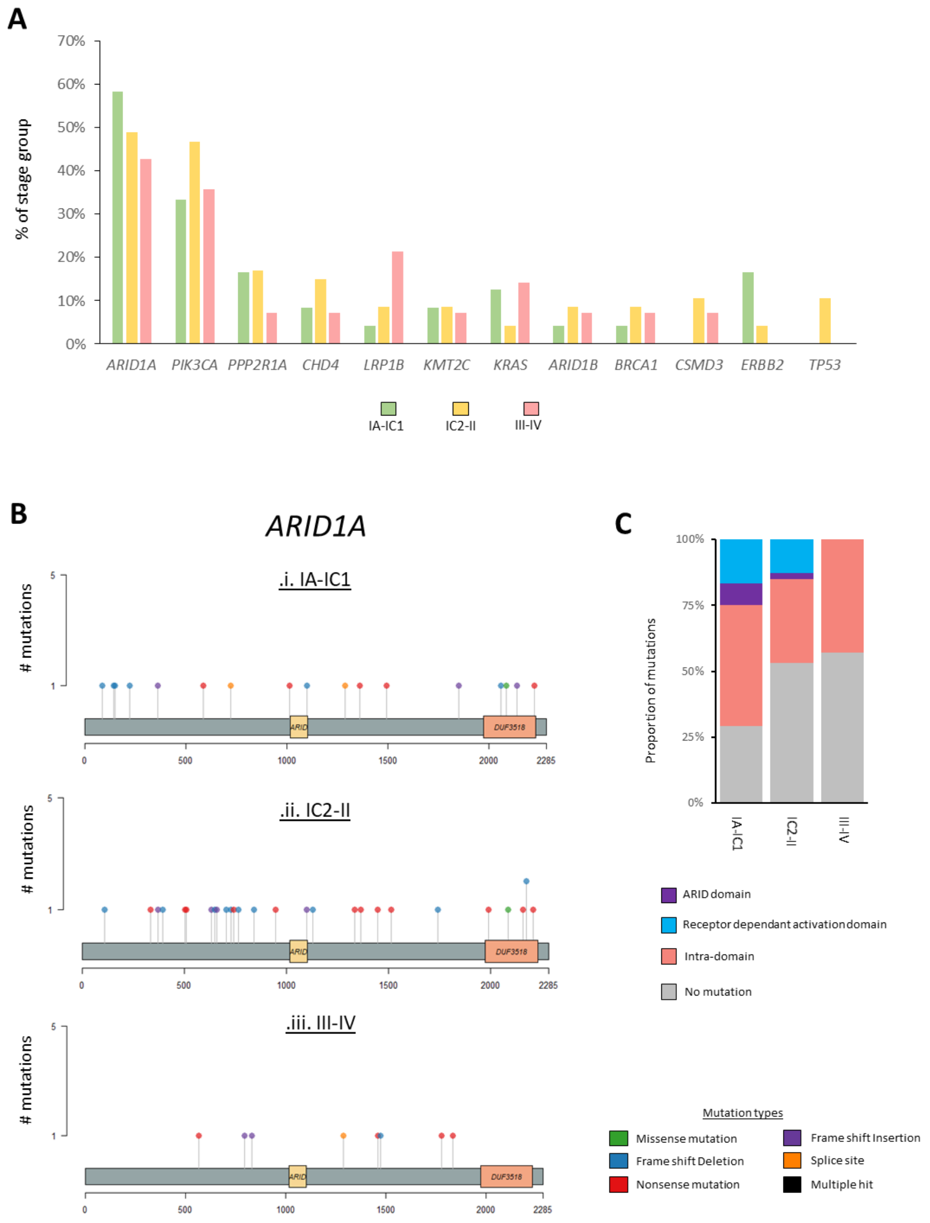


**Figure S5.** Mutations at frequently mutated genes in the cohort, split by stage group. **(A)** Bar plot of top 12 frequently occurring mutations, split by stage group, as a percentage of each group. **(B)** Lollipop plots of mutation types and positions within *ARID1A* split by stage group. Annotated domain regions are named. Key for mutation type is shown. **(C)** Proportional stacked plot of location of *ARID1A* mutations, split by stage group, plotted as a percentage

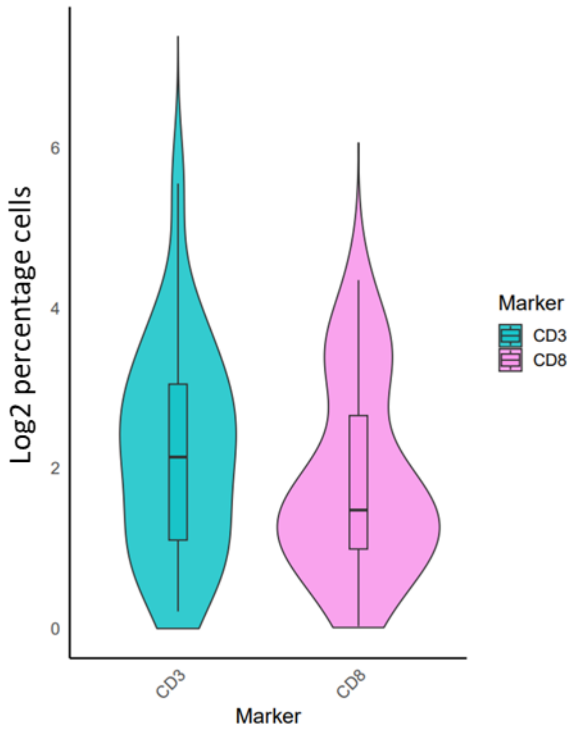


**Figure S6.** Violin plot of log2 percentage positive cells for either CD3 or CD8 markers. Boxes represent the interquartile ranges (IQR), with the middle lines indicating the median values. The whiskers extend to the highest and lowest data points that fall within 1.5 times the IQR.

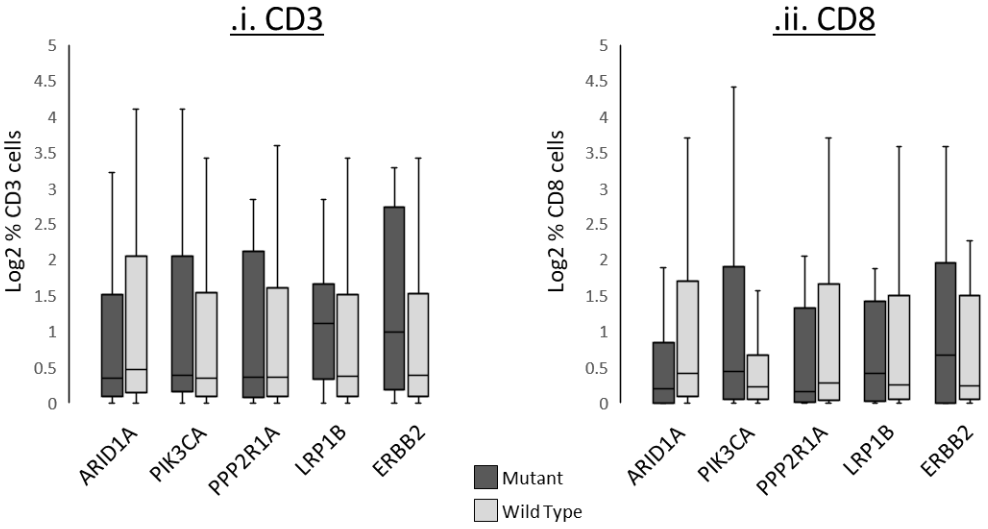


**Figure S7.** Box plot of log2 percentage positive cells for either CD3 or CD8 markers, split by mutational status of frequently mutated genes. Boxes represent the interquartile ranges (IQR), with the middle lines indicating the median values. The whiskers extend to the highest and lowest data points that fall within 1.5 times the IQR.

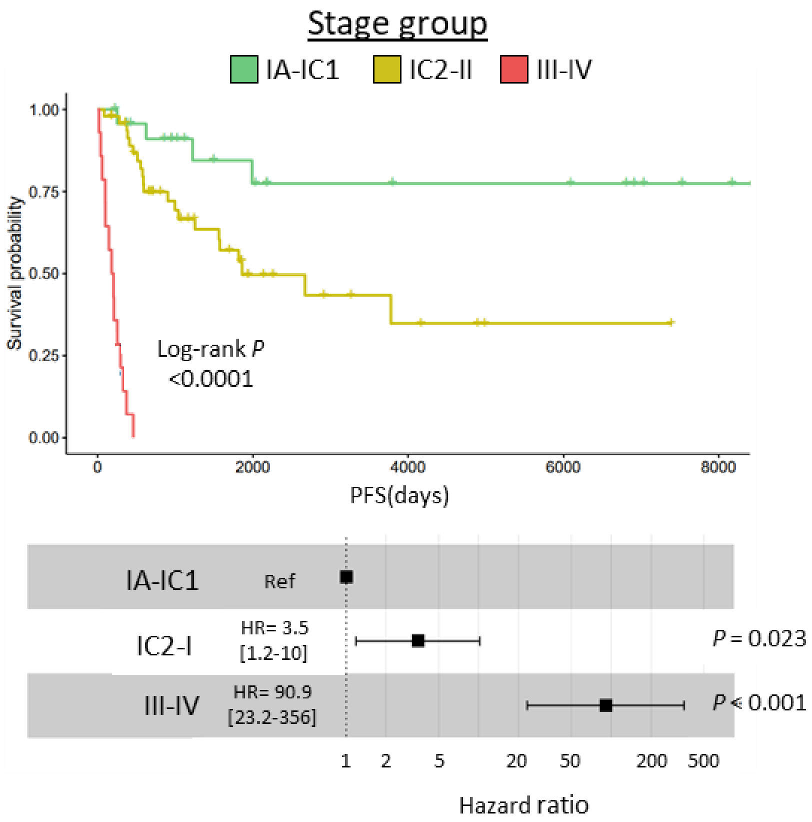


**Figure S8**. Kaplan-Meier survival analysis and hazard ratio plot for progression-free survival (PFS) in the study cohort when stratified by stage group. (Upper plots) Kaplan-Meier curve comparing PFS between groups, with statistical significance assessed by log-rank test. (Lower) Forest plot displaying the hazard ratio (HR) from a univariate Cox proportional hazards model for groups, with 95% confidence intervals.

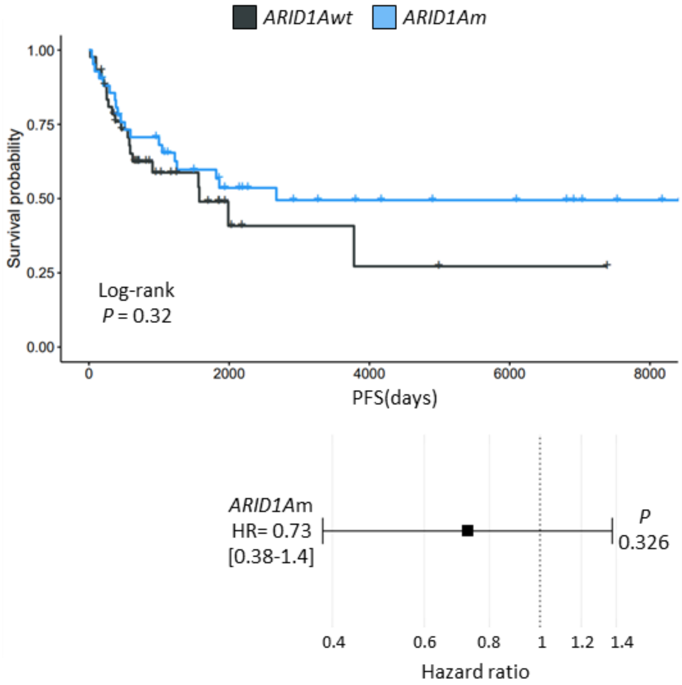


**Figure S9**. Kaplan-Meier survival analysis and hazard ratio plot for progression-free survival (PFS) in the study cohort when stratified by *ARID1A* mutation. (Upper) Kaplan-Meier curve comparing PFS between tumours containing wild type *ARID1A* (black) or mutated *ARID1A* (blue) tumours, with statistical significance assessed by log-rank test. (Lower) Forest plot displaying the hazard ratio (HR) from a univariate Cox proportional hazards model for mutated ARID1A, with 95% confidence intervals.

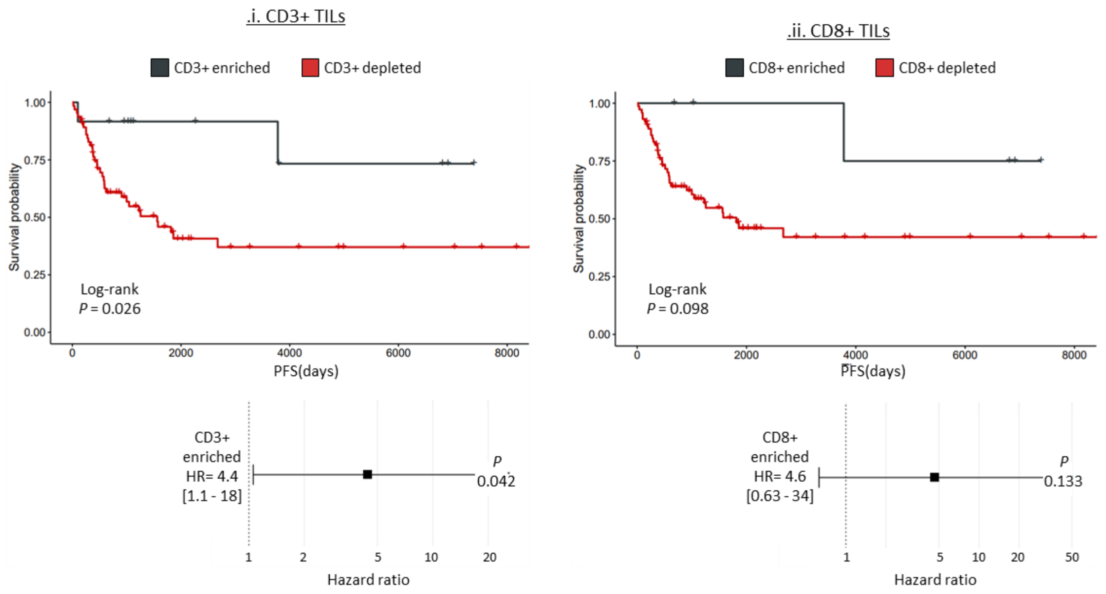


**Figure S10**. Kaplan-Meier survival analysis and hazard ratio plot for progression-free survival (PFS) in the study cohort when stratified by (i) CD3+ or (ii) CD8+ tumour infiltrating lymphocyte (TIL) levels. (Upper plots) Kaplan-Meier curve comparing PFS between TIL enrichment or deletion groups, with statistical significance assessed by log-rank test. (Lower) Forest plot displaying the hazard ratio (HR) from a univariate Cox proportional hazards model for groups, with 95% confidence intervals.

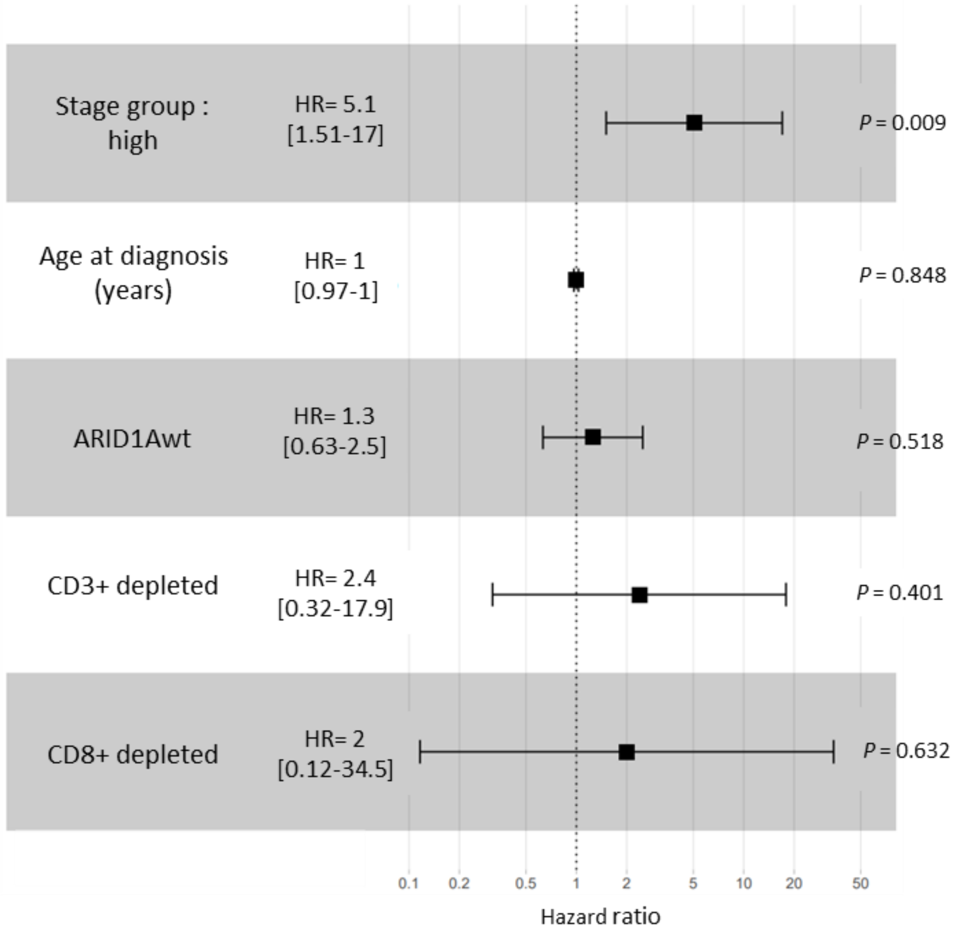


**Figure S11**. Multivariate Cox proportional hazards analysis for progression-free survival (PFS). Forest plot displaying hazard ratios (HR) with 95% confidence intervals for stage group "high", age at diagnosis (years), ARID1A wild-type status (ARID1Awt), low CD3+ TILs, and low CD8+ TILs, adjusted for covariates. P values are shown on the right.

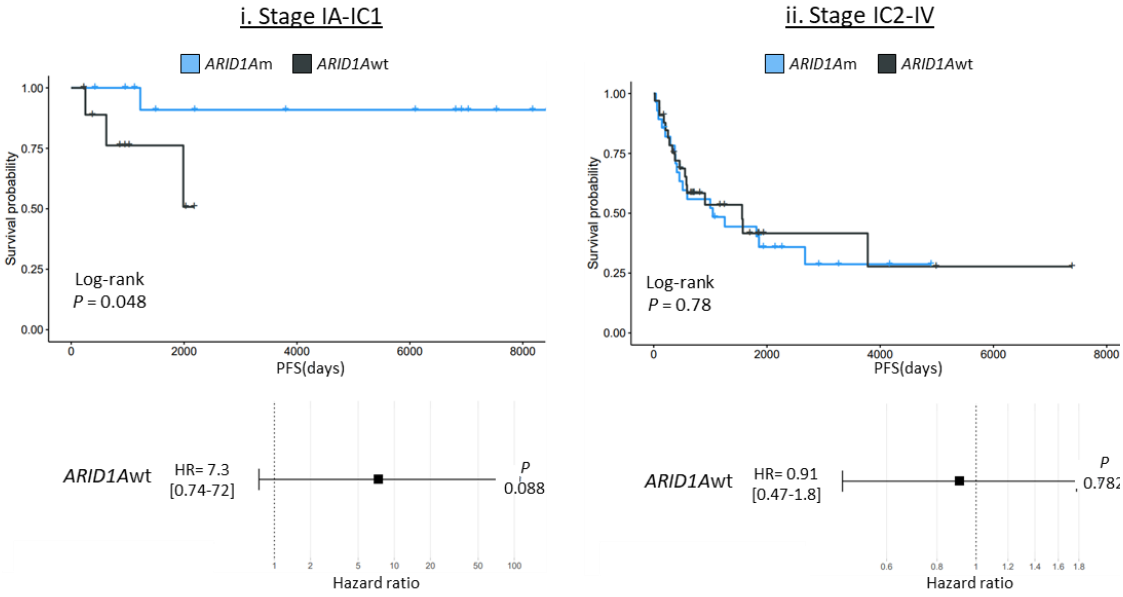


**Figure S12**. Kaplan-Meier survival analysis and hazard ratio plot for progression-free survival (PFS) in the study cohort when stratified by *ARID1A* mutational status for (i) low-stage IA-IC1 or (ii) high stages (IC2-IV). (Upper plots) Kaplan-Meier curve comparing PFS between TIL enrichment or deletion groups, with statistical significance assessed by log-rank test. (Lower) Forest plot displaying the hazard ratio (HR) from a univariate Cox proportional hazards model for groups, with 95% confidence intervals.

#### C. SUPPORTING TABLES

| | | | | |
|---|---|---|---|---|
| *AKT1* | *AKT2* | *ARID1A* | *AKT3* | *ARID1B* |
| *AGPAT9* | *AMER1* | *ATM* | *BAI3* | *BCL6* |
| *BRCA1* | *BRCA2* | *CCNE1* | *CHD4* | *CRHR1* |
| *CRKL* | *CTNNB1* | *CRTC1* | *CSMD3* | *DST* |
| *ERBB2* | *ETS1* | *FBXW7* | *KRAS* | *LIFR* |
| *KDM5C* | *KMT2C* | *KANSL1* | *LPHN3* | *LRP1B* |
| *MTOR* | *MSH6* | *MDM4* | *MET* | *MLH1* |
| *MSH2* | *MSH3* | *MLH3* | *NF1* | *NFE2L2* |
| *NUMA1* | *PAK3* | *PIK3CA* | *PIK3CG* | *PIK3R1* |
| *PIK3R2* | *PKHD1* | *PMS1* | *PMS2* | *PDE4DIP* |
| *PPP2R1A* | *PTCH1* | *PTEN* | *RICTOR* | *SMARCA4* |
| *SF3B1* | *STK11* | *TLR4* | *ST7* | *SPOP* |
| *SYNE1* | *TGM7* | *TP53* | *TSC1* | *TSC2* |
| *UBAP1* | *WAS* | *ZNF217* | | |

**Supporting Table S1** The 68 genes targeted on the custom sequencing panel
